# ETV7 regulates breast cancer stem-like cell plasticity by repressing IFN-response genes

**DOI:** 10.1101/2020.09.02.279133

**Authors:** Laura Pezzè, Mattia Forcato, Stefano Pontalti, Kalina Aleksandra Badowska, Dario Rizzotto, Ira-Ida Skvortsova, Silvio Bicciato, Yari Ciribilli

## Abstract

Cancer stem cells (CSCs) represent a population of cells within the tumor able to drive tumorigenesis and known to be highly resistant to conventional chemotherapy and radiotherapy. In this work, we show a new role for ETV7, a transcriptional repressor member of the ETS family, in promoting breast cancer stem-like cells plasticity and resistance to chemo- and radiotherapy in breast cancer (BC) cells. We observed that MCF7 and T47D BC-derived cells stably over-expressing ETV7 showed reduced sensitivity to the chemotherapeutic drug 5-Flouororuacil and to radiotherapy, accompanied by an adaptive proliferative behavior observed in different culture conditions. We further noticed that alteration of ETV7 expression could significantly affect the population of breast CSCs, measured by CD44^+^/CD24^low^ cell population and mammosphere formation efficiency. By transcriptome profiling, we identified a signature of Interferon-responsive genes significantly repressed in cells over-expressing ETV7, which could be responsible for the increase in the breast CSCs population, as this could be partially reverted by the treatment with IFN-β. Lastly, we show that the expression of the IFN-responsive genes repressed by ETV7 could have prognostic value in breast cancer, as low expression of these genes was associated with a worse prognosis. Therefore, we propose a novel role for ETV7 in breast cancer stem cells’ plasticity and associated resistance to conventional chemotherapy and radiotherapy, which involves the repression of a group of IFN-responsive genes, potentially reversible upon IFN-β treatment. We, therefore, suggest that an in-depth investigation of this mechanism could lead to novel breast CSCs targeted therapies and to the improvement of combinatorial regimens, possibly involving the therapeutic use of IFN-β, with the aim of avoiding resistance development and relapse in breast cancer.

## INTRODUCTION

Breast cancer is the most frequent tumor and the leading cause of cancer-related deaths in women ^1^. Breast cancer is a highly genetically heterogenous disease ^2^, and current therapeutic approaches are chosen based on the subtype of cancer, stage, mass and localization of the tumors among several parameters ^3^. The most common therapeutic strategies rely both on local (e.g., surgery and radiotherapy) and systemic treatments (e.g., chemotherapy, endocrine/hormonal and targeted therapy) ^4^. Hormone-dependent breast cancers, such as luminal breast cancers, frequently become refractory to the initially effective hormonal treatments, thus eventually requiring chemotherapy ^5^. Chemotherapy, in fact, still represents the most common option for advanced breast cancers. However, it has been demonstrated that chemotherapy itself, despite triggering shrinkage of primary tumors, can enhance metastasis through induction of EMT ^6^. Moreover, cancer cells can develop drug resistance, resulting in treatment failure and recurrence ^7, 8^. Cancer stem cells (CSCs) are considered the population of cells within the tumor able to drive tumorigenesis. CSCs are characterized by self-renewal ability and differentiation potential, and can therefore contribute to the heterogenicity of cancer cells within the tumor ^9, 10^. Moreover, they are known to be intrinsically highly resistant to conventional anti-cancer agents, including chemotherapy and radiotherapy ^11, 12^. Therefore, uncovering the pathways and mechanisms involved in CSCs enrichment, development of drug resistance or other unwanted side effects associated with chemotherapy is an urgent and critical aim for cancer research, oriented to improve treatment efficacy.

Using genome-wide transcriptional profiling of MCF7 human breast adenocarcinoma-derived cells, we identified ETV7 as a potential tumor-promoting factor among the genes more synergistically up-regulated by the combinatorial treatment with the chemotherapeutic agent Doxorubicin and the inflammatory cytokine TNFα ^13^.

ETV7, also called TEL2 or TELB, is a member of the large family of ETS (E26 Transforming Specific) transcription factors, whose members can regulate the expression of genes involved in various processes, such as development, differentiation, cell proliferation, migration, and apoptosis ^14, 15^. Given their role in essential cellular functions, their dysregulation can result in severe impairments within the cell, as demonstrated by the involvement of ETS factors in various diseases ^16–18^. In particular, many ETS factors have been associated with cancer initiation, transformation and metastatic spread ^19–22^. The roles of ETV7 are still poorly understood and studied, partially because of the absence of ETV7 gene in most rodent species, including mice ^23^. However, several studies highlighted multiple roles for ETV7 in hematopoiesis. For examples, the over-expression of ETV7 in both human and mice hematopoietic stem cells (HSCs) could increase the proliferation and deplete HSCs ^24^. Moreover, ETV7 over-expression in human U937 cells was shown to impede monocytic differentiation ^25^. ETV7 is also recognized as an Interferon (IFN)-stimulated gene (ISG), as its expression was shown to be induced by type I, type II and type III IFN treatment in different cell types ^26–28^. Elevated ETV7 expression has been associated with several tumor types; however, the role of ETV7 in cancer has been poorly investigated. ETV7 was shown to cooperate with Eμ-MYC in promoting B-lymphomagenesis ^29^ and the forced expression of ETV7 in mouse bone marrow was shown to cause myeloproliferative diseases, even if with a long latency, suggesting TEL2 as a bona fide oncogene ^30^. Besides, the crossing of the latest established ETV7 transgenic mouse model with an established leukemic mouse model revealed a remarkable acceleration in Pten^-/-^ leukemogenesis ^31^. Further evidence supporting ETV7 pro-tumorigenic functions comes from the recent identification of a transcriptional-independent activity of ETV7, which was shown to physically interact with mTOR into the cytoplasm generating a novel complex called mTORC3, which contributes to resistance to rapamycin, an mTOR-targeting anti-cancer agent ^32^. In contrast with these observations, ETV7 was shown to act as tumor suppressor in nasopharyngeal carcinoma by repressing SERPINE1 ^33^, and its down-regulation was observed in drug-resistant cancer cells ^34^ Recently, Piggin and colleagues reported an average higher level of ETV7 expression in tissues from all the breast cancer subtypes compared to normal breast tissues, with a correlation of ETV7 expression and breast cancer aggressiveness ^35^. We have recently demonstrated in breast cancer-derived cells that ETV7 expression can be induced by DNA-damaging chemotherapy, and elevated ETV7 levels has been associated with reduced sensitivity to Doxorubicin treatment, a mechanism which involve ETV7-mediated direct repression of DNAJC15, resulting in the increase in ABCB1 expression levels ^36^.

In this study we further demonstrated the role of ETV7 in breast cancer cell resistance to another chemotherapeutic drug, 5-Fluorouracil (5-FU) and radiotherapy. Noteworthy, we were able to observe a remarkable increase in breast cancer stem cells content in the cells over-expressing ETV7. We finally identified via RNA-seq a set of type I and type II Interferon response genes that were repressed by ETV7 as putative responsible of the stem cells enrichment.

In the light of these results, we propose a novel role for ETV7 as a regulator of breast cancer stem cell-like plasticity, which is mediated by the repression of IFN-stimulated genes and that can be partially reverted by the stimulation of IFN response with IFN-β.

## MATERIALS AND METHODS

### Cell lines and culture conditions

MCF7 cells were obtained from Interlab Cell Line Collection bank (IRCCS Ospedale Policlinico San Martino, Genoa, Italy), T47D cells were received from Dr. U. Pfeffer (IRCCS Ospedale Policlinico San Martino), HEK293T cells were a gift from Prof. J. Borlak (Hanover Medical School, Germany), while MDA-MB-231 cells were a gift from Prof. A. Provenzani (CIBIO Department, University of Trento, Italy).

MCF7, T47D, and HEK293T cells were grown in DMEM medium (Gibco, ThermoFisher Scientific, Milan, Italy) supplemented with 10% FBS (Gibco), 2mM L-Glutamine (Gibco) and a mixture of 100U/ml Penicillin / 100μg/ml Streptomycin (Gibco). MDA-MB-231 cells were cultured in the same medium with the addition of 1% Non-Essential Amino acids (Gibco). Cells were grown at 37°C with 5% CO2 in a humidified atmosphere. Cell lines were monthly checked for mycoplasma contaminations and have recently been authenticated by PCR-single-locus-technology (Eurofins Genomics, Ebersberg, Germany or DDC Medical, Fairfield, OH, USA).

### Treatments

5-FluoroUracil (5-FU) (Sigma Aldrich, Milan, Italy) was used at different concentrations and for different time points based on the specific experiment. Recombinant human IFN-β and IFN-γ (Peprotech, Tebu-Bio, Milan, Italy) were used at different concentrations and for different time points based on the experiment. Radiation treatment was performed using the Elekta Precise Linear Accelerator (Elekta Oncology Systems, Crawley, UK) for different doses and time points.

### Plasmids

The expression vector pCMV6-Entry-ETV7 C-terminally tagged with DDK-Myc tags was purchased from Origene (Tema Ricerca, Bologna, Italy). The lentiviral vector pAIP-ETV7 was obtained by cloning using the following primers to amplify the *ETV7* gene from pCMV6-Entry-ETV7 and inserting it into the pAIP-Empty plasmid (the tails containing restriction endonucleases’ target sequences are indicated in lowercase):

Fw: aggttaacATGCAGGAGGGAGAATTGGCTA
Rv: gagaattcTTAAACCTTATCGTCGTCATCC

pAIP was a gift from Jeremy Luban (Addgene plasmid #74171; http://n2t.net/addgene:74171; Watertown, MA, USA). Purified PCR product was inserted into pAIP backbone using HpaI and EcoRI restriction endonucleases (New England Biolabs; Euroclone, Milan, Italy). Correct cloning was checked by restriction analysis and direct sequencing (Eurofins Genomics).

pCMV-D8.91 and pCMV-VSVg plasmids for lentiviral particles production were obtained from Prof. Anna Cereseto, Laboratory of Molecular Virology, CIBIO Department, University of Trento, Italy.

### Viral vectors production in HEK293T

To obtain viral particles, HEK293T packaging cells were seeded into P150 dishes and transiently transfected with a mix containing 17.5 μg pCMV-D8.91 plasmid, 7.5 μg pCMV-VSVg plasmid, 25 μg lentiviral vector containing the gene of interest and 112.5 μl PEI 2X transfection solution (Sigma-Aldrich). After 48 hours, viral vectors containing the plasmid of interest were collected in the supernatant and filtered through a 0.45 μm filter. Vectors yield was quantified by the SG-PERT (Product-Enhanced Reverse Transcriptase) assay with the help of Prof. Anna Cereseto’s group, Laboratory of Molecular Virology, CIBIO Department, University of Trento, Italy.

### Generation of stable pAIP-ETV7 and Empty MCF7 and T47D cells

To obtain cell lines having a stable over-expression of the *ETV7* gene, MCF7 and T47D cells were transduced with 1 RTU of the lentiviral vector pAIP-Empty and with the lentiviral vector for the expression of the heterologous gene *ETV7*, pAIP-ETV7. After 72 hours, cells were split, and Puromycin (Life Technologies) was added at a concentration of 1.5 and 2.5 μg/ml respectively for MCF7 and T47D cells. Each 3 days medium was replaced, and after 4 cycles of selection, single cell cloning was performed for MCF7 cells according to the Corning protocol for cell cloning by Serial dilution in 96-well plates. During the single cell cloning procedure Puromycin concentration was gradually reduced to 0.75 μg/ml. T47D did not undergo serial dilution but were kept as the pooled population under Puromycin selection (1.5 μg/ml).

### RNA isolation and RT-qPCR

Total RNA was extracted using the Illustra RNA spin Mini Kit (GE Healthcare, Milan, Italy), converted into cDNA with the RevertAid First Strand cDNA Synthesis Kit following manufacturer’s recommendations (ThermoFisher Scientific) and RT-qPCR was performed with 25 ng of template cDNA into 384-well plates (BioRad, Milan, Italy) using the Kapa Sybr Fast qPCR Master Mix (Kapa Biosystems, Resnova, Ancona, Italy) or the qPCRBIO SyGreen 2X (PCR Biosystems, Resnova) and the CFX384 Detection System (BioRad). *YWHAZ* and *GAPDH* were used as housekeeping genes to obtain the relative fold change with the ΔΔCt method as previously described ^37^. Primer sequences were designed using Primer-BLAST designing tool (https://www.ncbi.nlm.nih.gov/tools/primerblast/), checked for specificity and efficiency, and are listed in Supplementary Table 1 (Eurofins Genomics).

### Western Blot

Western blotting was performed as previously reported ^38, 39^. Briefly, total protein extracts were obtained by lysing the cells in RIPA buffer (150mM Sodium Chloride, 1% NP-40, 0.5% Sodium Deoxycholate, 0.1% SDS, and 50mM TrisHCl pH 8.0) and proteins were quantified with the BCA method (Pierce, ThermoFisher Scientific); 20-50 μg of protein extracts were loaded on appropriate 7.5%, 10% or 12% polyacrylamide gels and SDS-PAGE was performed. Proteins were then transferred on Nitrocellulose membranes, which were probed over-night at 4°C with specific antibodies diluted in 1% non-fat skim milk-PBS-T solution: ETV7/TEL2 (E-1, sc-374478), β-Tubulin (3F3-G2, sc-53140), STAT1 (D1K9Y, Cell Signaling Technology, Euroclone), p-STAT1 (Tyr701) (D4A7, Cell Signaling Technology), BCL-2 (100, sc-509), Survivin (D-8, sc-17779), GAPDH (6C5, sc-32233), and α-Actinin (H-2, sc-17829). Antibodies were obtained from Santa Cruz Biotechnologies (Milan, Italy) when not explicitly indicated. Detection was performed with ECL Select reagent (GE Healthcare) using the UVITec Alliance LD2 (UVITec Cambridge, UK) imaging system.

### Cell Titer Glo Viability Assay

Viability assay was performed using the Cell Titer-Glo Luminescent cell viability assay (Promega) according to the manufacturer’s instructions. Cells were seeded in white flat 96-well plates and treated with different concentrations of 5-FU for 72 hours. Afterward, the plates were equilibrated at room temperature for 30 minutes; then, 100 μl of Cell Titer-Glo reagent was added to 100 μl of medium and left in incubation for 2 minutes on an orbital shaker. Then, a luminescence measure was performed at the Infinite M200 plate reader (Tecan). Viability was calculated as a % ratio of viable cells treated with the indicated drug respect to DMSO control.

### Vi-CELL Viability Assay and doubling time calculation

The cell viability after radiotherapy treatment was measured using the Vi-CELL viability analyzer (Beckman Coulter), which performs trypan blue dye exclusion method with an automated system. Cells were seeded in 6-well plates for 24 hours and then treated with different radiation doses (2 to 10 Gy) and incubated for 72 hours. Afterward, cells were harvested using trypsin and re-suspended in PBS. Vi-CELL performed the automated count of viable cells in the sample, allowing for viability measurement. The viability was calculated as the percentage of viable cells respect to the untreated control. Doubling time was calculated with the following formula:

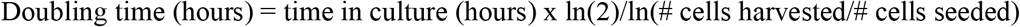

### Annexin V-FITC/PI staining

Apoptosis was measured using the FITC Annexin V Apoptosis Detection Kit I (BD Biosciences) as previously described ^40^. Briefly, cells were stained with an antibody against Annexin V conjugated to FITC fluorochrome together with the vital dye propidium iodide (PI). After the appropriate treatment, cells were harvested, washed twice with cold PBS and re-suspended in 1X Annexin V Binding Buffer at the concentration of 1.5×10^6^ cells/ml. 100 μl of the cell suspension was then incubated with 2.5 μl of FITC Annexin V and 5 μl of PI for 15 minutes at room temperature in the dark. Subsequently, 400 μl of 1X Annexin V Binding Buffer was added to each tube and samples were then analyzed. Flow cytometry analysis was conducted at the CIBIO Cell Analysis and Separation Core Facility using a FACS Canto A instrument (BD Biosciences).

### CD44/CD24 staining

The membrane expression of CD44 and CD24 cell surface markers was measured by double staining with antibodies conjugated with fluorophores and flow cytometry analysis. Cells were seeded in 6-well plate or T25 flasks and, after the appropriate treatments, were harvested and washed with PBS. 3×10^3^ cells were resuspended in 30 μl PBS + 0.1% BSA and incubated with APC mouse anti-human CD44 (cat.no 559942, BD Bioscience) and FITC mouse anti-human CD24 (cat.no 555427, BD Bioscience) antibodies or with their isotype controls (FITC mouse IgG2a, k isotype, and APC mouse IgG2b, k isotype, BD Bioscience) in ice for 30 minutes. After incubation cells were washed three times with PBS and finally re-suspended in 300 μl PBS. Flow cytometry analysis was performed as mentioned above.

### ALDEFLUOR analysis

ALDEFLUOR kit (STEMCELL Technologies, Cologne, Germany) was used to identify cells expressing high levels of ALDH enzyme. Experiments were performed following the manufacturer’s instructions. Briefly, cells were harvested and re-suspended in ALDEFLUOR Assay Buffer at the concentration of 3×10^5^ cells/ml. Then, 5 μl of activated ALDEFLUOR reagent was added to the cells and mixed. 500 μl of the cell suspension and reagent mixture was immediately moved to the control tube containing 5 μl of DEAB (diethylaminobenzaldehyde) reagent, a specific inhibitor of ALDH activity. Samples and controls were incubated for 45 minutes at 37°C. After the incubation, cells were centrifuged and re-suspended in ALDEFLUOR Assay Buffer. Flow cytometry analysis was performed at the Tyrolean Cancer Research Institute using a FACS Canto II instrument (BD Biosciences).

### Colony formation assays

The clonogenic assay was performed seeding single cell suspension with 2×10^3^ MCF7 or 4×10^3^ T47D cells in 6-well plates in complete growth medium. Cells were let grow for 3 weeks changing the medium twice a week. After 3 weeks, the cell colonies were gently washed with PBS and stained by the incubation with 0.1% Crystal Violet solution in 20% methanol for 20 minutes. Excess staining was removed, and colonies were washed twice in PBS. Image analysis was performed with the Image J software in order to calculate the percentage of well’s area occupied by colonies. To characterize the capability of transformed cells to grow independently from a solid surface (anchorage-independent growth), we performed soft agar colony formation assay. The wells of 6-well plates were prepared with a layer of base agar (0.7% agarose in DMEM complete medium + 10% FBS). After the solidification of the base, single cell suspension containing 1.5×10^4^ (MCF7) or 2×10^4^ (T47D) cells/well was rapidly mixed to the soft agar solution (0.35% agarose in DMEM complete medium) and disposed of as a layer over the previously prepared base agar. Finally, 1 ml of growing medium was added on the top. Cells were let grow for 3 weeks changing the top medium twice a week. Afterward, cells were stained with of 0.1% Crystal Violet solution in 20% methanol and washed several times with PBS 1X to remove the excess of Crystal Violet solution. Image analysis was performed with the Image J software in order to calculate the average size of the colonies.

### Mammospheres culturing

To generate primary mammospheres, cells were harvested with trypsin, centrifuged and re-suspended in mammosphere medium (DMEM/F12 supplemented with 2mM L-Glutamine, 100U/ml penicillin, 100μg/ml streptomycin with the addition of 20ng/ml recombinant human Epidermal Growth Factor (EGF), 10ng/ml recombinant human basic Fibroblast Growth Factor (bFGF) and 1x B27 supplement (Life Technologies). Cells were then counted and passed through a 25G needle to obtain a single cell suspension. Then, 10^3^ cells/well were seeded in ultra-low attachment 24-well plates (Corning, Rome, Italy) in 800 μl of mammosphere medium. Plates were incubated at 37°C for one week, and images were obtained at DM IL LED Inverted Microsocope (Leica). Mammospheres were split by harvesting the mammospheres with PBS, centrifuging and re-suspending with TrypLE Express Reagent (ThermoFisher Scientific) as an alternative to trypsin. The single cell suspension was then again counted, and seeded at the same starting concentration in mammosphere medium. After 1 week, mammospheres were counted and it was possible to calculate the Mammosphere Forming Efficiency (%) using the following equation:

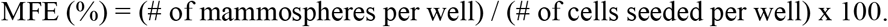

### Gene expression profiling

Total RNA from three independent biological replicates for MCF7 and T47D cells stably transduced with pAIP-Empty and pAIP-ETV7 was checked for quality and purity using the Agilent 2100 Bioanalyzer (Agilent Technologies, Santa Clara, CA, USA) selecting RNA extracts with a RIN (RNA integrity number) value > 8. RNA (1 μg) was then converted into cDNA libraries according to the Illumina TruSeq Stranded mRNA Sample Preparation Guide (Illumina, San Diego, CA, USA) following manufacturer’s instructions. Pair-end reads (2×75 bp) were generated from the libraries using the Illumina HiSeq2500 sequencer at CIBIO NGS Core Facility according to the standard Illumina protocol.

Raw reads were adapter-trimmed with BBDuk (sourceforge.net/projects/bbmap/) and aligned to build version hg38 of the human genome using STAR (version 2.5.3a; ^41^). Counts for UCSC annotated genes were calculated from the aligned reads using the *featureCounts* function of the *Rsubread* R package ^42^ and R (version 3.3.1). Normalization and differential expression analysis were carried out using *edgeR* R package ^43^. Raw counts were normalized to obtain Counts Per Million mapped reads (CPM). Only genes with a CPM greater than 1 in at least 3 samples were retained for downstream analysis. Gene expression changes were considered statistically significant with Benjamini-Hochberg FDR ≤5%. Enrichment analysis for Gene Ontology - Biological process category was performed on the list of 427 down-regulated genes upon ETV7 over-expression in both cell lines using *enrichGO* function of the *clusterProfiler* R package ^44^. Gene set enrichment analysis ^45^ was performed in preranked mode on the log2 fold change values estimated by edgeR and the Hallmark collection of Molecular Signature Database (MSigDB version 6.1, ^46^). Gene sets were considered significantly enriched at FDR ≤5% when using 1,000 permutations of the gene sets. We defined a signature of 22 ETV7-regulated IFN-responsive genes taking genes significantly down-regulated in both cell lines upon ETV7 over-expression and included in the HALLMARK_INTERFERON_ALPHA_RESPONSE and HALLMARK_INTERFERON_GAMMA_RESPONSE gene sets of MSigDB. Heatmaps were created using standardized CPM expression values and *pheatmap* R package (https://CRAN.R-project.org/package=pheatmap). RNA-seq data from this study have been deposited at Gene Expression Omnibus database (GEO, https://www.ncbi.nlm.nih.gov/geo/) with accession number GSE152580.

FPKM normalized expression data for TCGA breast cancer samples were downloaded from NCI Genomic Data Commons (TCGA-BRCA project) using the *TCGAbiolinks* R package ^47^. ETV7 expression in different sample types (i.e., primary tumor, metastasis, and normal tissue) and PAM50 molecular subtypes was compared using Analysis of variance (ANOVA) and post-hoc Tukey test. Scores for the ETV7-regulated IFN-responsive gene signature in primary tumor samples have been calculated as the average expression of the genes comprised in the signature. Samples have been classified as signature “high” if the ETV7-regulated IFN-responsive gene signature score was higher than the median of the signature scores in the tumor samples, and “low” vice versa. To evaluate the prognostic value of the ETV7-regulated IFN-responsive gene signature, we estimated, using the Kaplan-Meier method, the probabilities that patients would survive (overall survival, OS; disease specific survival, DSS), would remain free of disease (disease free interval, DFI) or progression (progression free interval, PFI). Survival data were censored at 12 years. Kaplan-Meier curves were compared using the logrank test in the *survcomp* R package.

### Statistical analysis

Where not specified, statistical analyses were performed using the GraphPad Prism version 6.0 software. When appropriate, unpaired t-test was applied for statistical significance. We selected throughout this study the two-sample Student’s t-test for unequal variance, given that we generally compared two conditions (i.e., treated vs. untreated samples, or over-expression of ETV7 vs. Empty control).

## RESULTS

### Increased ETV7 expression decreases the sensitivity of cancer cells to 5-FluoroUracil and to radiotherapy

We previously showed that ETV7 over-expression in breast cancer cell lines could reduce the sensitivity to the chemotherapeutic drug Doxorubicin ^36^. To test whether ETV7 altered expression could affect the sensitivity to other therapeutic treatments for breast cancer, we established two luminal breast cancer-derived cell lines (i.e., MCF7 and T47D) stably over-expressing ETV7 by lentiviral transduction of a plasmid carrying the ETV7 cDNA or the empty counterpart. The over-expression of ETV7 in MCF7 and T47D was verified by western blot analysis (Fig.1 A and B). We first evaluated the sensitivity of the cells to 5-FluoroUracil (5-FU), a commonly used breast cancer chemotherapeutic drug which, similarly to Doxorubicin, is also able to induce ETV7 expression in breast cancer cells ^36^. We analyzed the viability of the cells after treatment with 5-FU using Cell Titer Glo Assay, and we could observe that the cells over-expressing ETV7 showed a remarkable reduced sensitivity to the treatment compared to their empty counterpart, in both MCF7 (Figure 1C) and T47D cells (Figure 1D). To confirm the increased resistance to 5-FU of the cells over-expressing ETV7, we analyzed the induction of cell death, and in particular of apoptosis, in cells treated with 5-FU. By Annexin V-FITC/PI staining and flow cytometry analysis, we observed a significant reduction in PI positive (dead) cells (Figure 1E) and a significant, but less pronounced, reduction in the percentage of Annexin V positive (apoptotic) cells (Suppl. Fig. 1A). We previously demonstrated that the ETV7-induced resistance to Doxorubicin involved drug efflux mediated by the ABC transporter ABCB1, which was up-regulated at the transcriptional level both in MCF7 and MDA-MB-231 cells over-expressing ETV7 ^36^. We decided to extend our analysis to other ABC transporters, which are commonly deregulated in drug-resistant breast cancers, such as ABCC1/MRP1 and ABCG2/BCRP ^48–50^. We observed ABCB1, ABCC1, and ABCG2 up-regulation in MCF7 cells over-expressing ETV7 (Figure 1F); however, no significant differences could be observed in their expression in T47D cells (Suppl. Fig. 1B). Noteworthy, we were able to detect a remarkable increase in the expression of the anti-apoptotic proteins BCL-2 and Survivin in MCF7 cells over-expressing ETV7 (Figure 1G), which could explain the decreased percentage of apoptotic cells upon 5-FU treatment visible in MCF7 cells.

**Figure 1.**
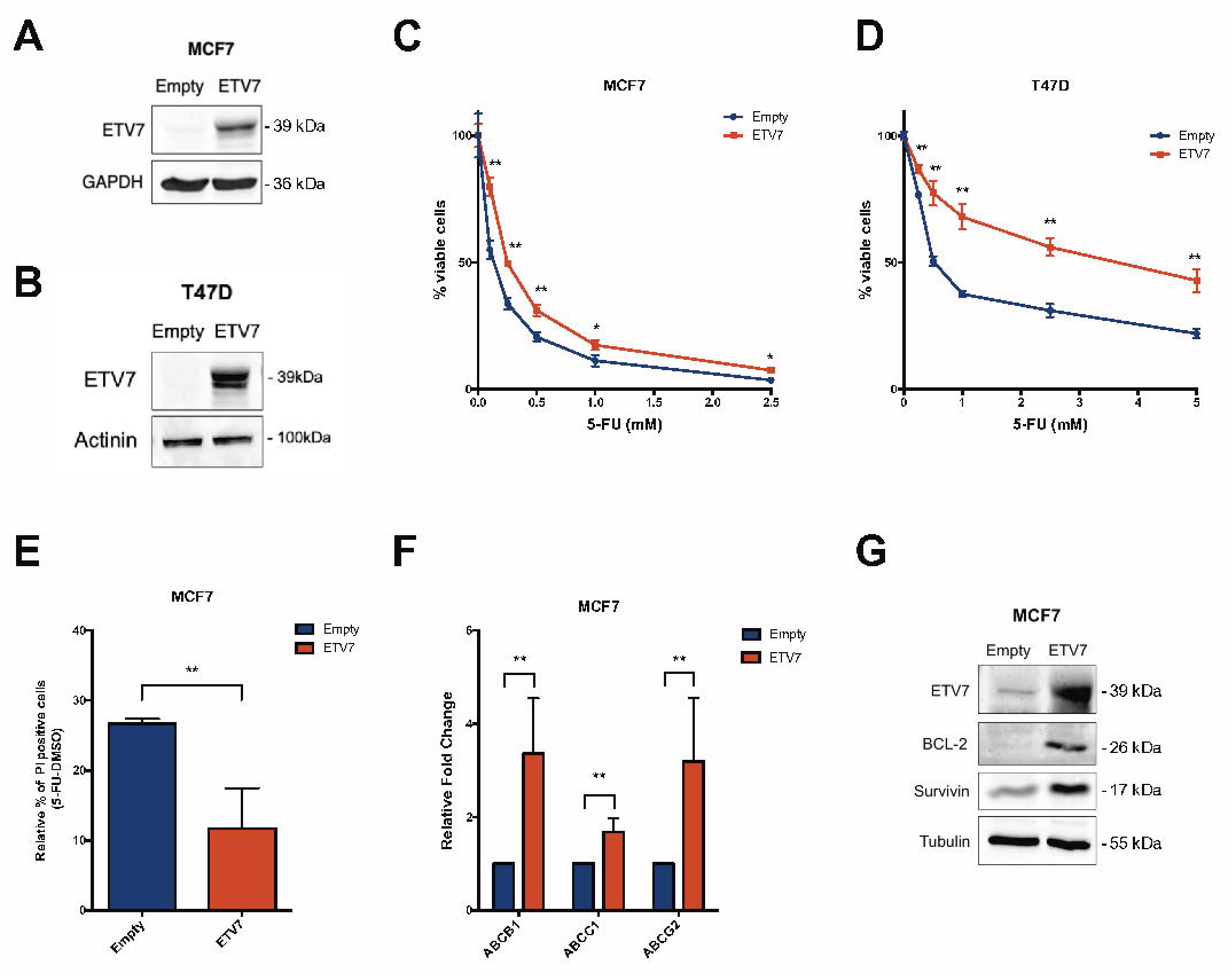
ETV7 expression modulates 5-FU sensitivity in breast cancer-derived MCF7 and T47D cells. A-B) Western blot analysis to measure ETV7 protein expression in MCF7 (A) and T47D (B) cells stably over-expressing ETV7 or their empty control. GAPDH, Actinin and Tubulin expression were used as loading control. Blots were cropped for clarity and conciseness of the presentation. C-D) Cell Titer Glo Assay for survival analysis upon 5-FU treatment in MCF7 (C) and T47D (D) cells over-expressing ETV7 and their empty control. The percentage of viable cells was calculated normalizing the luminescence measures on the DMSO treated sample. E) Relative percentage of PI positive cells calculated as the difference of 5-FU and DMSO treated cells measured by Annexin V-FITC/PI staining of MCF7 Empty and MCF7 ETV7 cells treated with 5-FU 200 μM for 72 hours.33 F) RT-qPCR analysis of ABCB1, ABCC1, and ABCG2 expression in MCF7 Empty and MCF7 ETV7 cells. G) Western Blot analysis of the anti-apoptotic BCL-2 and Survivin protein expression in MCF7 Empty and MCF7 ETV7 cells. Tubulin was used as loading control. Bars represent the averages and standard deviations of at least three independent experiments. On the left of each blot is indicated the approximate observed molecular weight. * = p-value < 0.05; ** = p-value < 0.01.

In order to test whether the over-expression of ETV7 could confer resistance to other breast cancer treatments that act through a different mechanism of action, we examined the sensitivity to radiotherapy. We exposed the cells to different doses of radiation (from 2 to 10 Gy) and measured the percentage of viable cells and the induction of apoptosis. Interestingly, cells over-expressing ETV7 showed a decreased sensitivity also to radiotherapy exposure in both MCF7 and T47D cells (Suppl. Fig. 1C and 1D). However, significant differences between ETV7 over-expressing cells and the empty control in the induction of apoptosis could be detected only for the lower dose of radiation (2Gy) in both the cell lines (Suppl. Fig. 1E e 1F), suggesting that the resistance mechanism induced by ETV7 relies only partially on the inhibition of apoptosis.

### The over-expression of ETV7 alters the proliferative potential of luminal breast cancer cells

Since the protective effect determined by ETV7 seems to be only partially dependent on the inhibition of apoptosis, we analyzed the proliferative potential of the cells; in fact, as both 5-FU and radiotherapy exert their action when the cells divide, their activity is strongly dependent on the proliferative rate of the cells. We first analyzed the proliferative potential of MCF7 and T47D cells stably over-expressing ETV7 by measuring their doubling time using the ViCell instrument. We observed a significant increase in the doubling time of MCF7 cells over-expressing ETV7 in comparison to MCF7 Empty cells, demonstrating that cell proliferation was slower in cells over-expressing ETV7 compared to the empty control (Suppl. Fig. 2A). In T47D cells, this increase in the doubling time was instead slight and not significant (Suppl. Fig. 2B). Using clonogenic assay, we could appreciate a substantial decrease in the colony formation potential of cells over-expressing ETV7 when grown on plastic in both MCF7 and T47D cells (Figure 2A and 2B). Therefore, the decreased proliferative rate of the cells could also explain the increased resistance to 5-FU and radiotherapy. Notably, despite the similar proliferation rate when cultured in normal conditions, the growth of T47D Empty cells was significantly reduced compared with MCF7 Empty cells when grown at single cell density. Surprisingly, the analysis of the colony formation ability of the cells in an anchorage-independent system with the soft agar assay showed an opposite proliferative behavior. Indeed, both MCF7 and T47D cells over-expressing ETV7 formed larger colonies when grown in soft agar compared to control (Figure 2C and 2D). Nevertheless, the anchorage-independent growth is commonly considered a pro-tumorigenic feature; thus, these apparently contradictory results might suggest that the cells over-expressing ETV7 could gain the ability to switch to different proliferative behaviors based on the environmental conditions.

**Figure 2.**
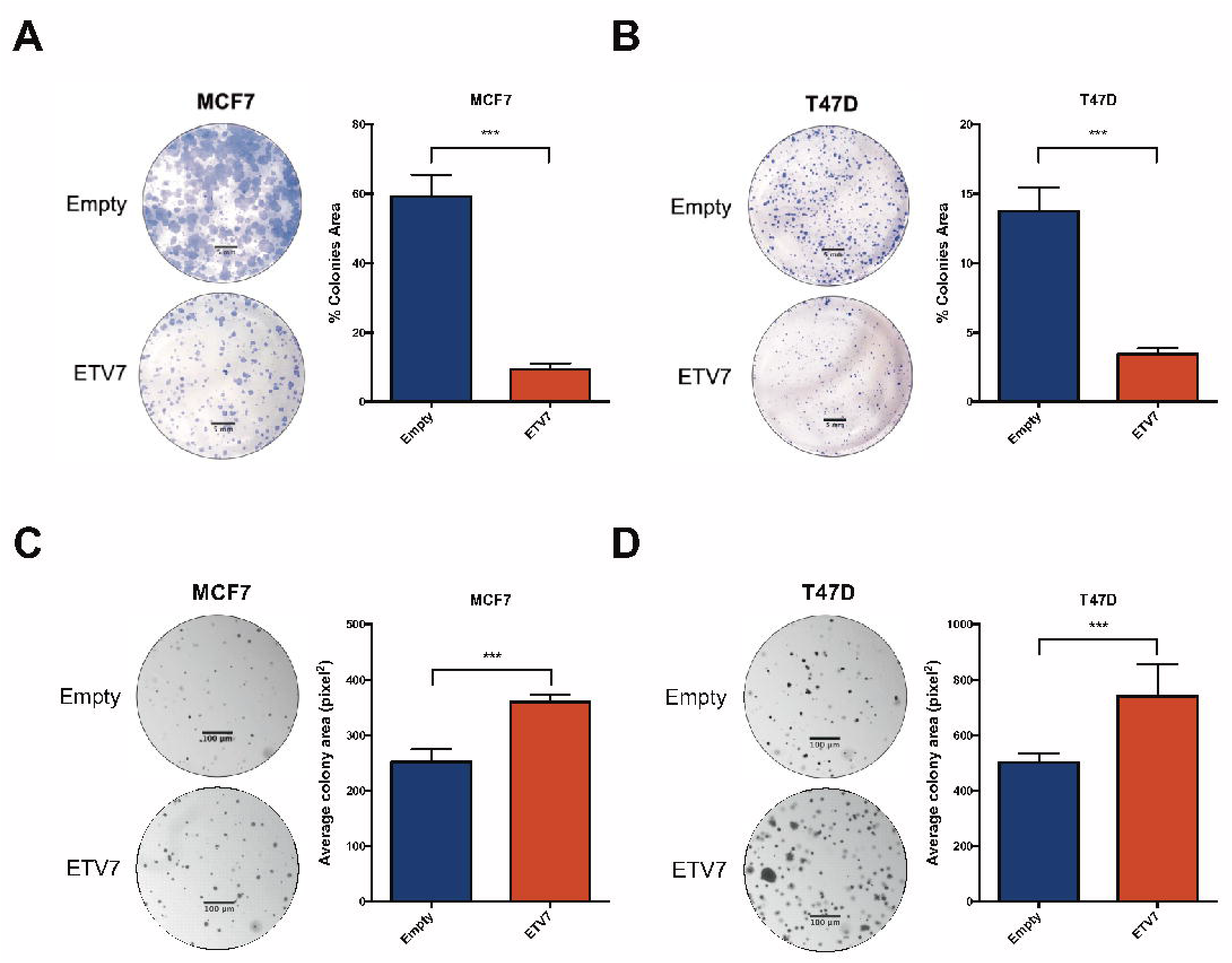
Breast cancer-derived cells over-expressing ETV7 display altered proliferative potential. A-B) Colony formation/clonogenic assay of MCF7 (A) and T47D (B) Empty and ETV7 cells after 3 weeks of growth. Presented are a representative image (left) of the entire well and the total colony area (right), calculated with the ImageJ software. C-D) Soft agar colony formation assay of MCF7 (C) and T47D (D) Empty and ETV7 cells after 3 weeks of growth. On the left, a representative image of part of the well (taken at 5X magnification) and on the right the average colony area calculated with the ImageJ software. Indicated are the scale bars for microscopy images. Bars represent the averages and standard deviations of at least three independent experiment. *** = p-value < 0.001.

### The expression of ETV7 affects BCSCs markers expression and mammosphere formation

Given the observed increased resistance to 5-FU and radiotherapy, the adaptive proliferative potential of the cells to different environment conditions, and data from the literature reporting the involvement of ETV7 in cell differentiation ^25, 51^, we hypothesized that ETV7 might play a role in breast cancer stem-like cells plasticity. To test this hypothesis, we first analyzed some of the most commonly accepted markers for breast cancer stem cells, including CD44 and CD24 expression and ALDH activity. We measured the percentage of CD44^+^ and CD24^-^ cells (representing the BCSCs population) in MCF7 and T47D cells over-expressing ETV7 or their relative control. Both the parental cell lines presented a very low or almost absent population of CD44^+^/CD24^-^ cells; however, we could appreciate an impressive increase in this population in both the cell lines tested upon ETV7 over-expression (Figure 3A and 3B). Importantly, the over-expression of ETV7 could stimulate both the increase in CD44 expression and the decrease in CD24 expression on the plasma membrane of the cells. We then evaluated the activity of ALDH via ALDEFLUOR assay kit. Here again, we observed a very low population of ALDH+ cells in the parental cell lines, but, either a decrease in this already small population or no significant differences were observed in response to ETV7 over-expression in MCF7 (Suppl. Figure 3A) and T47D (Suppl. Figure 3B) cells, respectively. Thus, the over-expression of ETV7 seems to drive a strong polarization of the analyzed breast cancer cells toward a more cancer stem-like cell phenotype represented by the CD44^+^/CD24^-^ population.

**Figure 3.**
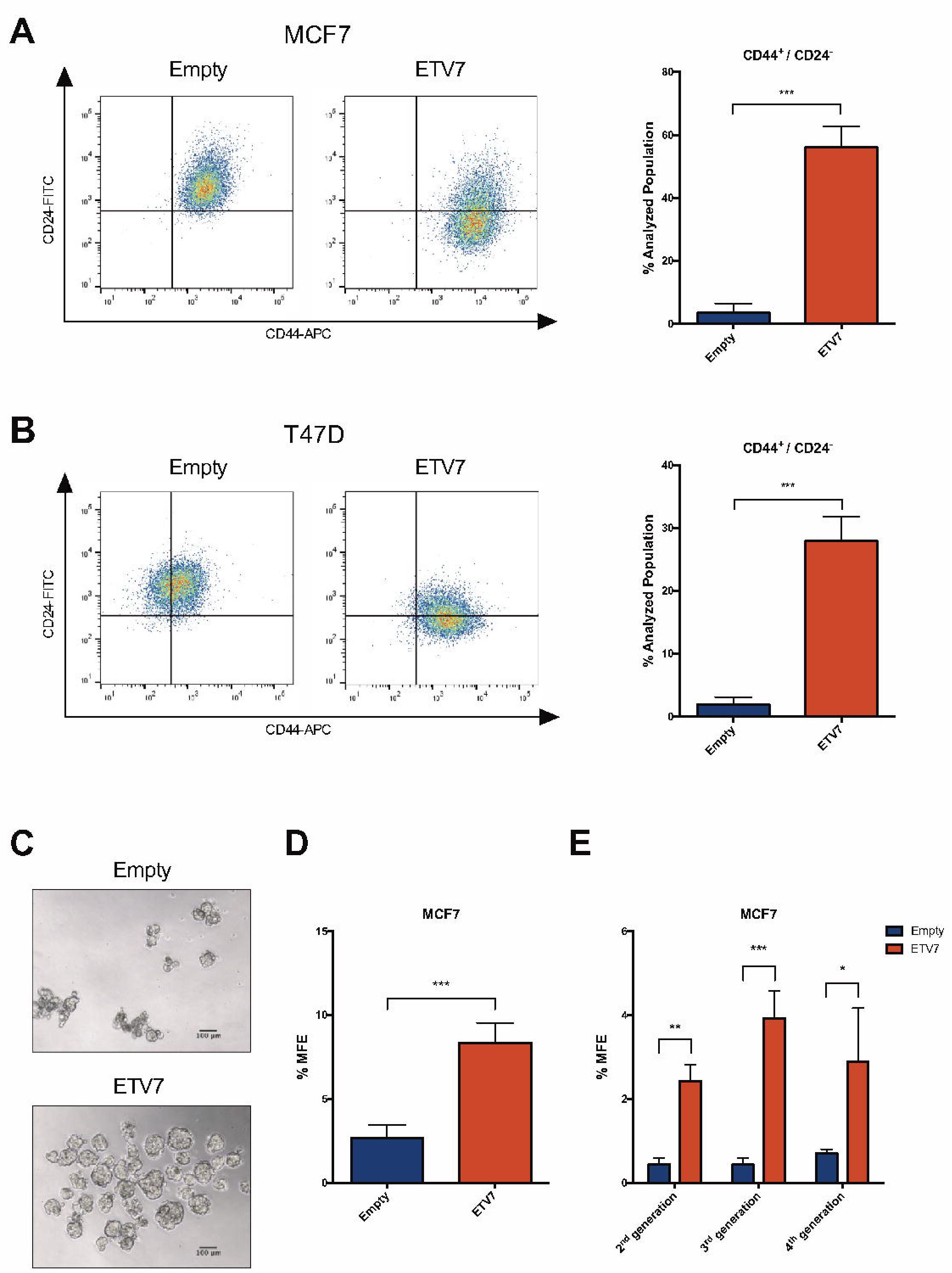
The over-expression of ETV7 influences the expression of breast cancer stem cell markers and mammosphere formation efficiency. A-B) CD44-APC and CD24-FITC staining and flow cytometry analysis in MCF7 (A) and T47D (B) Empty and ETV7 cells. On the left, a representative dotplot of the results obtained at FACS Canto A; the histogram on the right summarizes the percentage of CD44+/CD24-cells in Empty and ETV7 over-expressing cells. C) A representative image of first-generation mammospheres obtained from MCF7 Empty and MCF7 ETV7 cells, respectively. The scale bar is indicated. D) The percentage of mammosphere formation efficiency (% MFE) in MCF7 Empty and ETV7 calculated as number of mammospheres per well/number of cells seeded per well X 100. E) % MFE in second-, third-, and fourth-generation mammospheres obtained by passaging the mammospheres every 7 days. Bars represent the averages and standard deviations of at least three independent experiments. * = p-value < 0.05; ** = p-value < 0.01; *** = p-value < 0.001.

To verify whether modulating the expression of ETV7 in the opposite direction could also affect the population of CD44^+^/CD24^-^ cells, we silenced the expression of ETV7 in the aggressive triple negative BC-derived cell line MDA-MB-231, which is known from the literature to present almost exclusively CD44^+^/CD24^-^ cells ^52^. We knocked-down ETV7 expression using two different siRNAs (siETV7#1 and siETV7#2), and we confirmed their activity by RT-qPCR analysis (Suppl. Figure 3C). Noteworthy, although MDA-MB-231 cells do not express high level of ETV7, which does not account for their CD44/CD24 expression levels, in both samples silenced for ETV7 we observed a population of cells with decreased CD44 membrane expression compared to the scramble control, suggesting that tuning the expression of ETV7 could bi-directionally affect the population of CD44^+^/CD24^-^ cells (Suppl. Fig. 3D). Moreover, another feature of breast cancer stem cells is their potential to form spheres (i.e., mammospheres) when grown in non-differentiating and non-adherent conditions. We measured the mammosphere formation efficiency (MFE) of MCF7 cells over-expressing ETV7 and the relative empty control cells, and we noticed a significant increase in the MFE of MCF7 cells over-expressing ETV7 compared to the control (Figure 3C and Figure 3D). We then tested the propagation potential of mammospheres by passaging them in culture, and we could appreciate the ability of cells over-expressing ETV7 to generate second, third, and fourth generation mammospheres (Figure 3E). These data confirm that the over-expression of ETV7 in breast cancer cells can enhance the cancer stem-like cell properties and thus suggest a role for ETV7 in breast cancer stem-like cell plasticity.

Unfortunately, we could not obtain any mammosphere from T47D, which, when grown in non-adherent and non-differentiating conditions, could form only aggregates of cells with unorganized structure (data not shown).

### ETV7 induces the repression of a signature of interferon responsive genes

In order to identify the molecular mediators of the biological effects caused by the over-expression of ETV7, we performed a transcriptome analysis on MCF7 and T47D cells over-expressing ETV7 or the relative empty plasmid by RNA-seq. Indeed, being ETV7 a transcription factor, we expect it to exert most of its effects via transcriptional control. The analysis of transcriptional data identified 5,387 genes differentially expressed (DEGs) in MCF7 and T47D cells over-expressing ETV7 as compared to empty controls, respectively (Figure 4A). For each cell line, it was possible to observe a similar number of up- and down-regulated genes, with 721 genes which were commonly regulated in the two cellular systems (427 commonly down-regulated and 294 commonly up-regulated genes), and a lower number of genes which were inversely regulated in MCF7 and T47D cells (Figure 4A).

**Figure 4.**
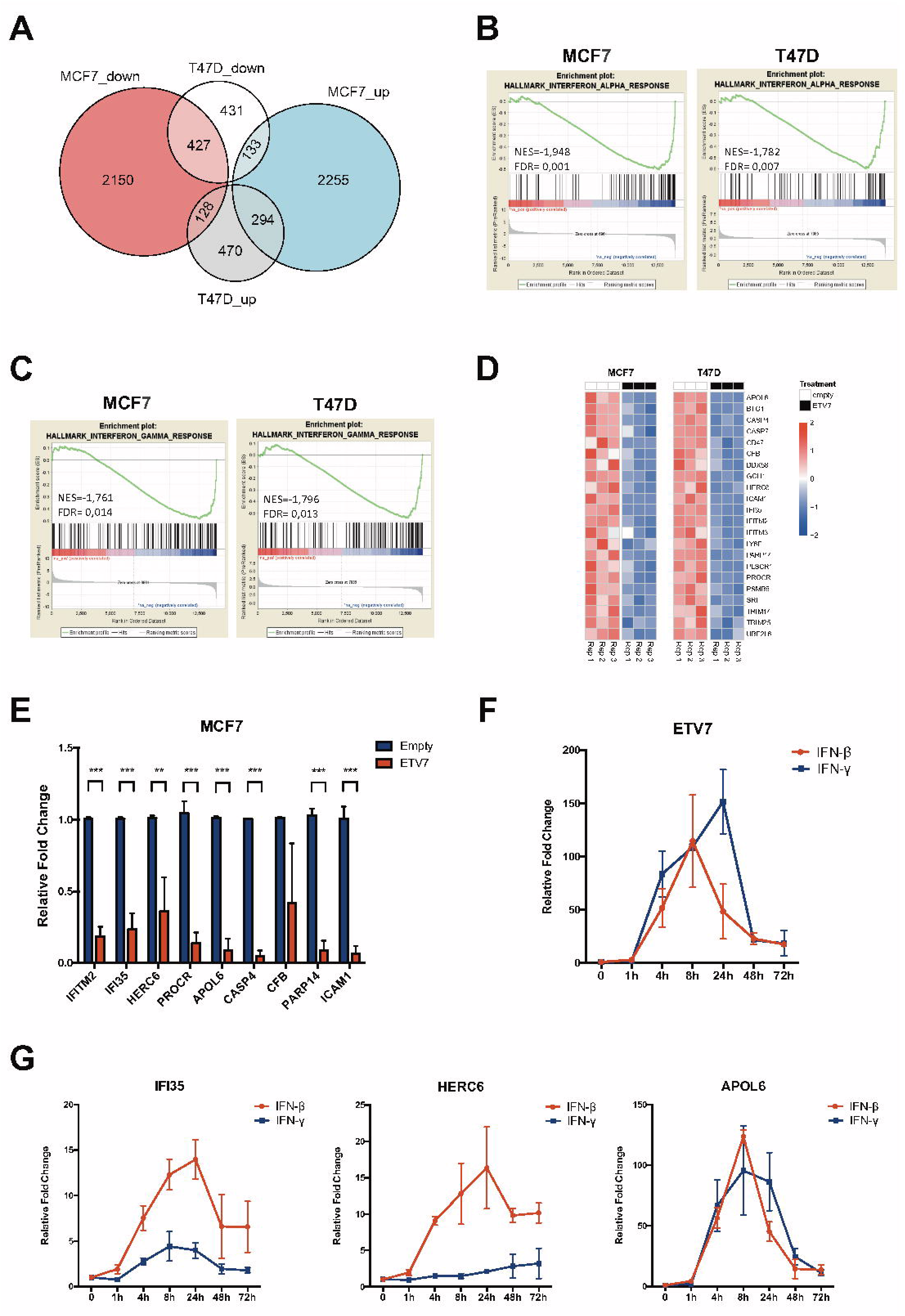
The over-expression of ETV7 causes the repression of a signature of IFN-response genes. A) A Venn diagram showing the number of differentially expressed genes (DEGs at a False Discovery Rate (FDR) ≤ 0.05) in the comparison between MCF7 and T47D cells over-expressing EVT7 and their respective controls. B-C) Gene Set Enrichment Analysis (GSEA) of MCF7 and T47D cells over-expressing ETV7 vs MCF7 and T47D controls. Enrichment plots for Type 1 Interferon response (B) and Type 2 Interferon response (C) gene sets of the Hallmark Collection. The Normalized Enrichment Score (NES) represents the degree of the enrichment of the gene set; the negative sign indicates that the gene set is down-regulated in cells over-expressing ETV7. D) Heatmaps showing the standardized expression level of genes comprising the ETV7-regulated IFN-responsive gene signature in MCF7 (left) and T47D (right) cells. E) RT-qPCR experiments for the validation of genes of the ETV7-regulated IFN-responsive signature with a Fold Change (FC) < −2 in MCF7 Empty and ETV7 cells. Bars represent the averages and standard deviations of at least three biological replicates. ** = p-value < 0.01; *** = p-value < 0.001. F) RT-qPCR analysis of normalized ETV7 expression relative to untreated control in MCF7 cells treated with 5ng/ml IFN-β (red) or IFN-γ (blue) at different time points. G) RT-qPCR analysis of the normalized expression of the genes regulated by ETV7 (IFI35, HERC6, APOL6) in MCF7 cells treated with 5ng/ml IFN-β (red) or IFN-γ (blue) at different time points. Bars represent the averages and SEM of at least three biological replicates.

Given that most of the biological effects previously observed in cells over-expressing ETV7 were common to both the cell lines, we focused our subsequent analyses on the common DEGs. In particular, as ETV7 is known to be a transcriptional repressor, we were interested in the commonly down-regulated genes, which could be regulated by ETV7 in a more direct way. Interestingly, the most significant terms obtained by the gene ontology analysis of common down-regulated DEGs involved innate immune response and inflammatory response, with several terms referred to the pathogen/viral entry into the host (Suppl. Fig. 4A). Moreover, gene set enrichment analysis (GSEA) highlighted “Interferon_alpha_response” (Figure 4B) and “Interferon_gamma_response” (Figure 4C), i.e., the Hallmark gene sets comprising genes involved in the cellular response to type I or type II Interferons, respectively, as the only common significantly repressed gene sets enriched in cells over-expressing ETV7. From these analyses, we then obtained a signature of Interferon responsive genes significantly down-regulated (FDR ≤ 0.05) in both the cell lines upon ETV7 over-expression (ETV7-regulated IFN-responsive gene signature; Figure 4D). We selected for validation the genes of the signature whose Fold Change was lower than −2 in both the cell lines, and we were able to confirm by RT-qPCR the repression of the entire list in MCF7 (Figure 4E) and T47D (Suppl. Fig. 4B) cells over-expressing ETV7, with the sole exception of CFB, whose significant repression could be validated only in T47D cells. We then decided to test the responsiveness of the selected genes to type I and type II IFN treatment in MCF7 cells. In particular, we chose to use IFN-β as type I IFN, given the previous data in literature suggesting a negative role for IFN-β in breast cancer stemness ^53, 54^, and IFN-γ as type II IFN, being the only member of this subfamily. Since, as mentioned above, ETV7 itself is a well-recognized Interferon-stimulated gene (ISG) ^26, 28^, we first analyzed its expression in response to IFN-β and IFN-γ in MCF7 cells. As expected, we observed a robust and significant induction of ETV7 expression in response to both types of IFNs, with a peak of induction at 8 hours for IFN-β and 24 hours for IFN-γ, confirming that ETV7 is strongly responsive to IFNs also in the breast cancer-derived MCF7 cell line (Figure 4F).

We then analyzed the expression of the validated genes repressed by ETV7 in response to IFN-β and IFN-γ. Interestingly, all the analyzed genes, with the exception of *ICAM1*, were more responsive to IFN-β than to IFN-γ (Figure 4G and Suppl. Fig. 4C).

### IFN-β treatment partially reverses the ETV7-dependent breast cancer stem-like cells plasticity

Given the observation that the over-expression of ETV7 could modulate breast cancer stem-like cells content in both MCF7 and T47D cells, and that this effect was reflected by a strong repression of a signature of IFN-responsive genes at the transcriptional level, we hypothesized that, if the repression of the IFN-responsive genes was responsible for the observed biological effects, the stimulation of their expression by treatment with IFNs could possibly reverse the breast cancer stem-like cell plasticity related to ETV7 expression. Therefore, we performed long-term treatments of MCF7 Empty and ETV7 cells with either IFN-β or IFN-γ and analyzed the mammosphere formation efficiency of the cells. Strikingly, IFN-β, but not IFN-γ, could strongly inhibit the mammosphere formation capacity of MCF7 cells over-expressing ETV7 already at the first generation (Figure 5A). Moreover, IFN-β treatment completely abolished the ability of the cells to propagate in vitro, as MCF7 cells over-expressing ETV7 could not generate second, third and fourth generation mammospheres (Figure 5B and Suppl. Fig. 5A and 5B). Our hypothesis was further supported by the analysis of CD44^+^/CD24^-^ cell population in MCF7 and T47D cells treated with either IFN-β or IFN-γ for 2 weeks. Indeed, treatment with IFN-β, but not with IFN-γ, could revert the increase in CD44^+^/CD24^-^ cells observed in the cells over-expressing ETV7 (Figure 5C and 5D). These data suggest that the over-expression of ETV7 could drive breast cancer stem-like cells plasticity through the repression of a panel of IFN-responsive genes, as the treatment with IFN-β, to which most of the validated genes are responsive, could reverse the breast cancer stem cells enrichment. This concept is further supported by the observation that, even in MCF7 cells over-expressing ETV7 where a strong repression in the expression of the IFN-responsive genes was visible, most of these genes were still responsive to the treatment (Suppl. Fig. 5C).

**Figure 5.**
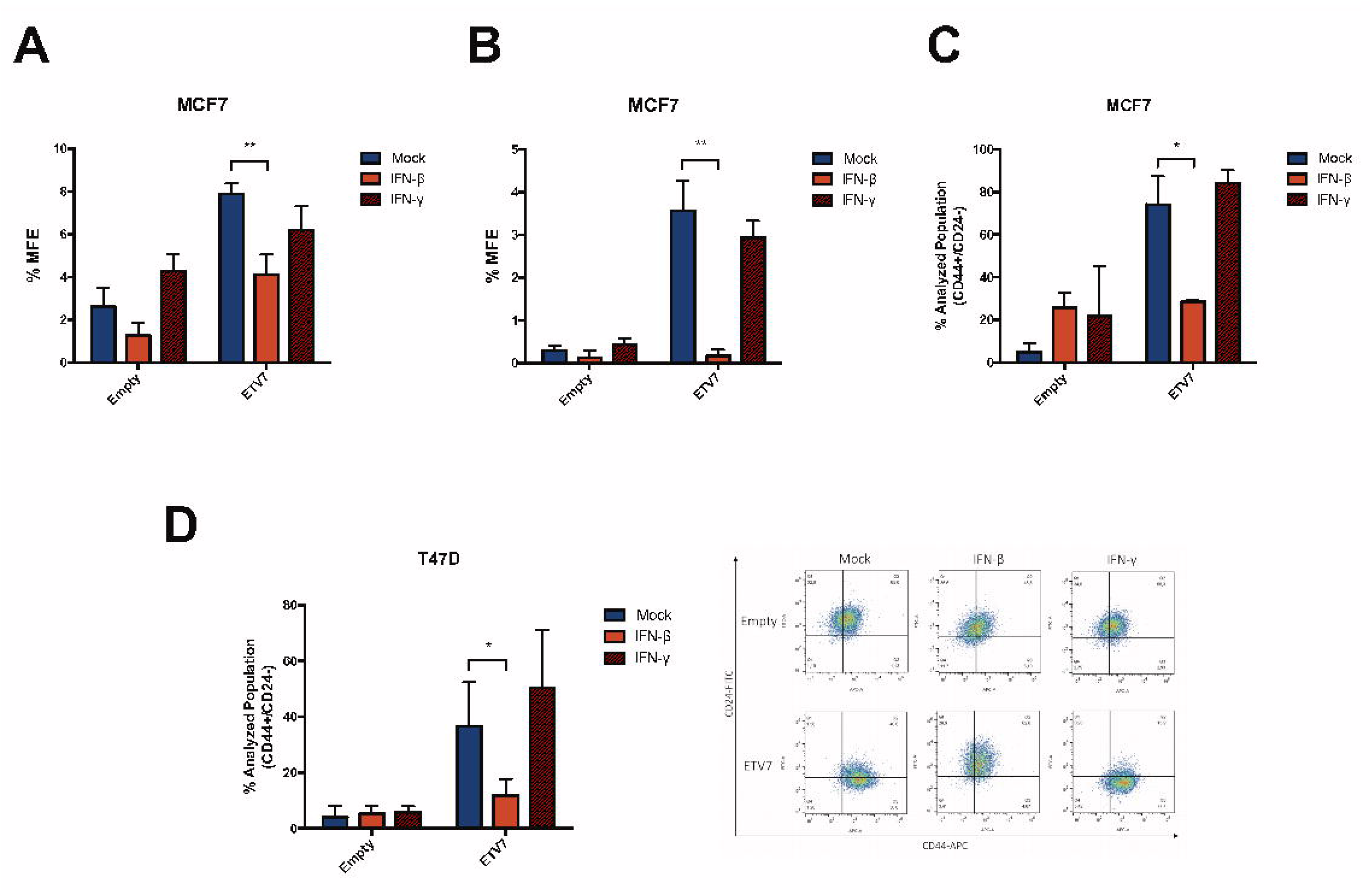
IFN-β treatment reverses the breast cancer stem-like cells enrichment induced by ETV7. A-B) Percentage of first (A) and second (B) generation mammosphere formation efficiency (% MFE) in MCF7 Empty and ETV7 cells in response to 5ng/ml IFN-β or IFN-γ calculated as number of mammospheres per well/number of cells seeded per well X 100. C-D) Histograms summarizing the percentage of CD44^+^/CD24^-^ cells measured by CD44-APC and CD24-FITC staining and flow cytometry analysis in MCF7 Empty and ETV7 cells (C) or T47D Empty and ETV7 cells (D) treated with 5ng/ml IFN-β and IFN-γ for 2 weeks. A representative dot plot of the results obtained at FACS Canto A in T47D cells is shown on the right. Bars represent the averages and standard deviations of at least three biological replicates. * = p-value < 0.05; ** = p-value < 0.01.

### ETV7-dependent repression of IFN-responsive genes shows prognostic value in breast cancer

To test whether the repression of the IFN-responsive genes expression mediated by ETV7 could be endowed with a prognostic significance in breast cancer patients, we first analyzed the expression of ETV7 in the breast cancer samples of the TCGA project (see methods for technical details). In accordance with a previous report ^35^, we could confirm a significant higher expression of ETV7 in metastatic and primary solid tumors as compared to normal tissue (Figure 6A). Moreover, the expression of ETV7 increased according to cancer aggressiveness, and it was significantly higher in basal-like samples compared to the other molecular subtypes (Figure 6B). Based on these findings, we then examined whether the expression of ETV7-repressed IFN-responsive gene signature could have a prognostic value in breast cancer patients (TCGA cohort). Noteworthy, Kaplan-Meier plots revealed that the low expression of the signature was significantly correlated with a lower probability of disease-specific survival (DSS, Figure 6C), overall survival (OS, Figure 6D), disease-free interval (DFI, Figure 6E) and progression-free interval (PFI, Figure 6F). We may speculate that the ETV7-dependent repression of the IFN-responsive genes could drive a more aggressive form of breast cancer, which according to our preliminary data, could possibly be responsive to IFN-β treatment.

**Figure 6.**
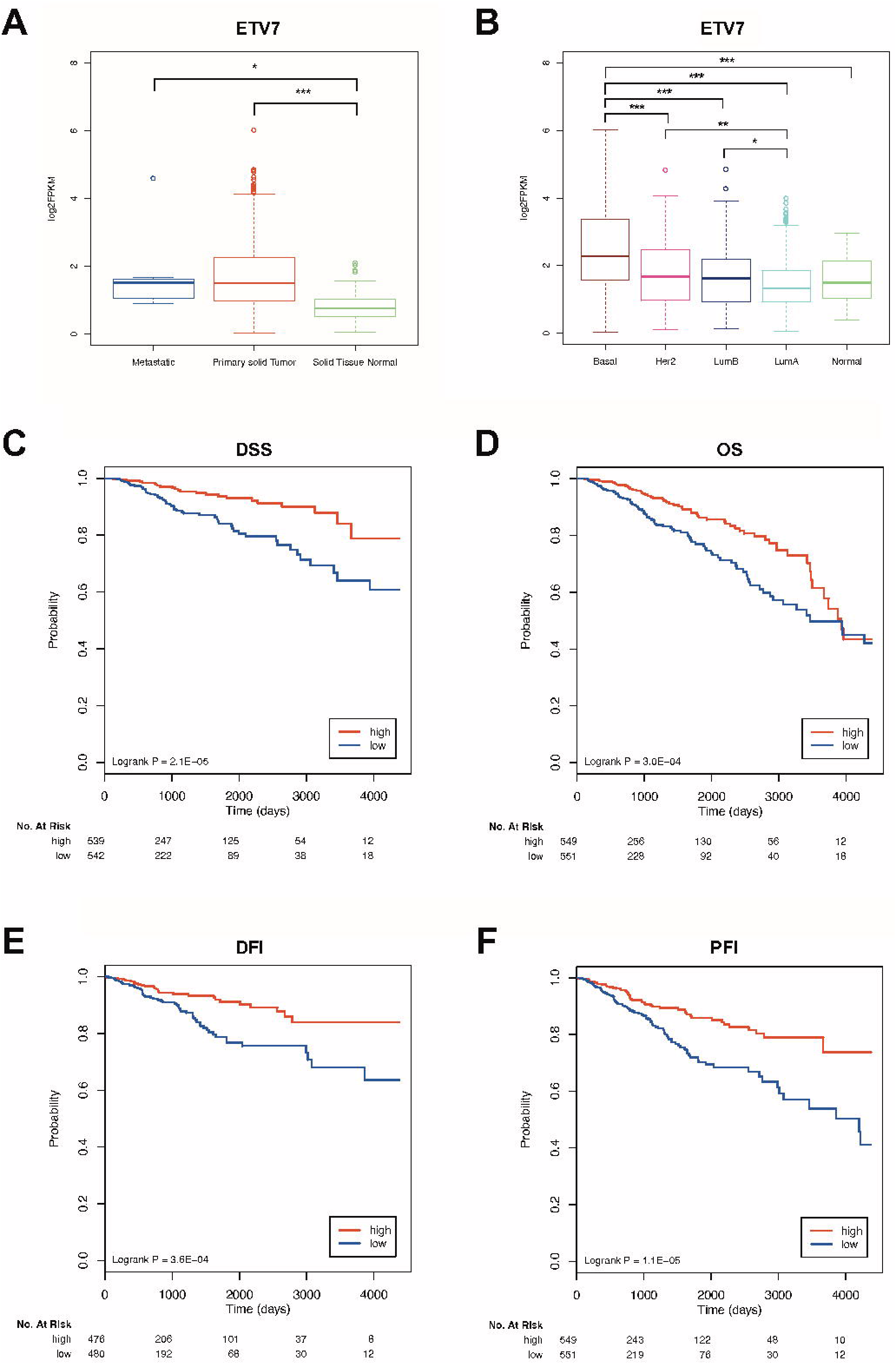
The expression of ETV7 and ETV7-repressed interferon signature genes have prognostic value in breast cancer. A, B) ETV7 expression in TCGA BRCA samples classified by tissue type (A) and PAM50 molecular subtype (B). Plot whiskers extend to the most extreme data point which is no more than 1.5 times the interquartile range from the box. Significance is calculated with ANOVA and post-hoc Tukey test. * = p-value < 0.05; *** = p-value < 0.001. C-F) Kaplan–Meier curves for TCGA breast cancer patients stratified according to the average expression of ETV7-regulated IFN-responsive gene signature. Curves represent the probability of disease specific survival (DSS) (C), overall survival (OS) (D), disease free interval (DSI) (E) and progression free interval (PFI) (F). p-values are calculated with logrank test.

## DISCUSSION

ETV7 is a poorly studied transcriptional repressor of the ETS family of transcription factors, which is mainly considered as an oncoprotein ^24, 30^. Recently, it was shown that the expression of ETV7 is significantly higher in breast cancer tissues compared to normal breast ^35^, suggesting a possible role for ETV7 in breast cancer pathogenesis; however, the functions and impact of ETV7 expression in breast cancer are still to be elucidated. We have recently published a novel mechanism of ETV7-mediated resistance to the chemotherapeutic agent Doxorubicin which involves the down-regulation of DNAJC15 tumor suppressor leading to an increased expression of ABCB1 efflux pump ^36^.

In this work, we expanded the effects of enhanced ETV7 expression on breast cancer progression and resistance to conventional anti-cancer drugs. Indeed, we demonstrated that the over-expression of ETV7 in MCF7 and T47D cells was also associated with reduced sensitivity to other types of standard treatments, such as 5-FU (Figure 1C-E and Supplementary Figure 1A) and radiotherapy (Supplementary Figure 1C-F). This increased resistance was accompanied by the induction of some ABC transporters frequently responsible for chemoresistance in breast cancer (i.e., ABCB1, ABCC1, and ABCG2) (Figure 1F and Supplementary Figure 1B), the increased expression of the anti-apoptotic proteins BCL-2 and Survivin (Figure 1G), and by the decreased proliferative potential observed in ETV7 over-expressing cells, since both 5-FU and radiotherapy efficacy depend on cell division (Figure 2A-B). Indeed, the observed increase in the expression of ABC transporters and anti-apoptotic proteins, together with the decreased cell proliferation, are common drug resistance-associated properties ^55–57^. Interestingly, we noticed that the proliferation rate of the cells underwent a switch when the cells were grown in an anchorage-independent manner. In particular, the colony formation potential in soft agar was significantly higher in cells over-expressing ETV7, whereas an opposite effect was observed when the colony formation efficiency was measured on plastic-support (Figure 2A-B vs. Figure 2C-D). This observation suggests a plastic behavior of the cells over-expressing ETV7, which can adapt their proliferative potential to the environmental conditions. Moreover, anchorage-independent growth is considered a pro-tumorigenic feature ^58^, suggesting that the over-expression of ETV7 may induce pro-tumorigenic effects in breast cancer cells. Given these observations, and the literature data reporting anti-differentiation roles for ETV7 ^29^, we hypothesized that the altered expression of ETV7 could affect the breast cancer stem cells population. Noteworthy, when we analyzed the CD44^+^/CD24^low/-^ BCSCs subpopulation in MCF7 and T47D cells over-expressing ETV7, we detected a remarkable increase (from ~ 1% in cells transfected with the empty vector to > 30% in both cell lines over-expressing ETV7) (Figure 3). This fascinating finding could be explained by the fact that CSCs can also originate from differentiated non-stem cells, through de-differentiation of normal somatic cells acquiring stem-like characteristics and malignant behavior via genetic or heterotypic alterations, such as EMT ^59^. Another recognized marker for BCSCs is the aldehyde dehydrogenase (ALDH), a family of cytosolic enzymes responsible for the detoxification by oxidation of intracellular aldehydes and are involved in the retinol to retinoic acid oxidation during stem cells differentiation ^60^. Surprisingly, using ALDEFLUOR assay, we found that MCF7 and T47D over-expressing ETV7 had a reduced or unchanged ALDH activity, respectively (Supplementary Figure 3A-B), in contrast with what mentioned above. However, CD44^+^/CD24^low/-^ BCSCs can overlap only partially with ALDH^+^ BCSCs, and present different properties. Liu and colleagues demonstrated that BCSCs can exist in two distinct states: a mesenchymal-like and an epithelial-like state ^61^. According to their work, CD44^+^/CD24^low/-^ cells belong to the mesenchymal-like BCSCs, are primarily quiescent, highly invasive, and are localized at the tumor invasive front. Conversely, the ALDH^+^ cells are the epithelial-like BCSCs, are highly proliferative, characterized by the expression of epithelial markers, and are localized more centrally within the tumor mass ^61^. This model can somehow explain the contradictory data about CSCs and the EMT state, as some studies suggest a commonality between the two states ^62^, whereas others suggest that these processes are mutually exclusive ^63^, and this is further proof of cancer stem cell plasticity in BC, as BCSCs can switch between the two states. In order to investigate genome-wide the mechanisms responsible for the effects of ETV7 observed in culture, we performed RNA-seq analysis on MCF7 and T47D cells over-expressing ETV7 and the relative controls. Since ETV7 is known to act as a transcriptional repressor, we focused our analysis mainly on the commonly down-regulated genes and gene sets in MCF7 and T47D cells over-expressing ETV7. Interestingly, the enrichment analysis revealed significant repression of a signature of Interferon-stimulated genes (ISGs) (Figure 4B-D and Supplementary Figure 4A). Therefore, we hypothesized that, if ETV7 exerts its effect on breast cancer stem cell plasticity via the repression of IFN response genes, inducing the re-expression of these genes might possibly reverse the observed effects. We thus selected IFN-β and IFN-γ as representative of type I and type II IFN, respectively, and tested the responsiveness of the obtained signature of ISGs in MCF7 cells. Interestingly, despite the fact that several genes were classified as type II ISGs in the MSigDB Hallmark collection, almost all of the analyzed genes were more responsive to IFN-β than to IFN-γ (Figure 4G and Supplementary Figure 4C). This effect can suggest the presence of a novel negative feedback loop, given that ETV7 itself is induced in response to IFNs as we also confirmed by RT-qPCR (Figure 4F). However, ETV7 expression did not affect STAT1 induction in response to IFN treatment, suggesting that the repression of the ISGs may act independently, or possibly downstream, on STAT1 activation (Supplementary Figure 6). Interestingly, we showed that the prolonged treatment of the cells with IFN-β, but not with IFN-γ, was able to rescue the effects on CSCs content tested both as CD44^+^/CD24^low/-^ population (Figure 5C-D) and by mammospheres formation ability (Figure 5A-B and Supplementary Figure 5A-B). The fact that IFN treatment, and in particular IFN-β, but not IFN-γ, can reverse the observed effects on CSC plasticity, supports the hypothesis that ETV7 mediates its functions via the repression of ISGs, as IFN-β is a stronger inducer of the ETV7-repressed genes compared to IFN-γ. Our results support the previous observations, which showed that immune-repressed triple-negative breast cancers (TNBC) lacking endogenous IFN signaling were highly recurrent, therapy-resistant, and characterized by CSC-like features ^64, 65^. Moreover, Doherty and colleagues recently showed that the treatment with IFN-β, which was able to restore the IFN signaling, could revert the CSCs properties observed in transformed mammary epithelial cells, suggesting it as a potential therapeutic approach for TNBC treatment ^53^. Our data strengthen these observations and provide a novel role for ETV7 in breast cancer stem-like cell plasticity. Lastly, we identified a gene signature of ETV7 down-regulated genes endowed of prognostic relevance in breast cancer patients, suggesting a role for ETV7 and for this set of genes as possible biomarkers for BC prognosis (Figure 6).

Taken collectively, our data reveal that the altered expression of ETV7 can affect the sensitivity of BC cells to some anti-cancer agents (i.e., Doxorubicin, 5-FU, and radiotherapy) and that it may accomplish this task by modulating the breast cancer stem cells plasticity. In light of these results, we can speculate that targeting CSCs in BC patients with IFNs (particularly IFN-β) prior (or in combination with) the administration of chemo/radiotherapy could enhance the efficacy of anti-cancer standard treatments.

## ACKNOWLEDGEMENTS

We thank CIBIO Next Generation Sequencing and CIBIO Cell Analysis and Separation Facilities for technical assistance. We are also thankful to Dr. U. Pfeffer, Prof. J. Borlak, and Prof. A. Provenzani for sharing cell lines and to Prof. A. Cereseto group for their help with viral infections and SG-PERT assay. We appreciated Prof. Alberto Inga and Dr. Alessandra Bisio for sharing reagents and helpful discussions. This work was supported by CIBIO Institutional Start-up funds (to YC) and by PRIN 2017 grant of the Italian Ministry of Education, University and Research (to MF; No. 2017HWTP2K_005).

---

**Suppl. Fig. 1.** A) A representative dotplot of the flow cytometry analysis performed on MCF7 Empty and MCF7 ETV7 cells treated with 5-FU 200 μM for 72 hours and the relative percentage of Annexin V positive cells calculated as the difference of 5-FU and DMSO treated cells. B) RT-qPCR analysis of ABCB1, ABCC1, and ABCG2 expression in T47D Empty and T47D ETV7 cells. C-D) ViCell Assay for survival analysis upon radiotherapy treatment in MCF7 (A) and T47D (B) cells over-expressing ETV7 and their empty control. E-F) Annexin V-FITC/PI staining of MCF7 (E) and T47D (F) cells Empty or over-expressing ETV7 treated with radiotherapy with 2-6-10 Gy for 72 hours. The relative percentage of Annexin V positive cells was calculated as the difference between treated and untreated cells. Bars represent the averages and standard deviations of at least four independent experiments. * = p-value < 0.05; ** = p-value < 0.01.

**Suppl. Fig. 2.** A-B) Doubling time of MCF7 (A) and T47D (B) Empty and ETV7 cells calculated by cell count at ViCell instrument. Bars represent the averages and standard deviations of at least four independent experiments. ** = p-value < 0.01.

**Suppl. Fig. 3.** A-B) ALDEFLUOR analysis in MCF7 (C) and T47D (D) Empty and ETV7 cells. The histogram on the left summarizes the percentage of ALDH positive cells in Empty and ETV7 over-expressing cells; on the right, a representative dot plot of the results obtained at FACS Canto II. C) RT-qPCR analysis of ETV7 expression in MDA-MB-231 transfected with siRNA against ETV7 (siETV7 #1 and #2) and the relative scramble control for 72 hours. Bars represent the averages and standard deviations of at least three independent experiments. * = p-value < 0.05; *** = p-value < 0.001. D) Representative dot plot of CD44-APC and CD24-FITC staining and flow cytometry analysis in MDA-MB-231 cells transfected with siRNA against ETV7 (siETV7 #1 and #2) and the relative scramble control for 72 hours.

**Suppl. Fig. 4.** A) Gene ontology analysis of commonly down-regulated DEGs in MCF7 and T47D cells (ETV7 vs. Empty). The number and fraction of commonly down-regulated DEGs that are annotated with a specific GO category are indicated by the dot size and on the x-axis, respectively. Dots are color-coded based on the enrichment adjusted p-values. In the image are shown the top 10 significant terms. B) RT-qPCR for validation of genes of the ETV7-regulated IFN-responsive signature with a Fold Change (FC) < −2 in T47D Empty and ETV7 cells. C) RT-qPCR analysis of the normalized expression of the genes regulated by ETV7 (IFITM2, CASP4, CFB, ICAM1, PARP14, PROCR) in MCF7 cells treated with 5ng/ml IFN-β (red) or IFN-γ (blue) at different time points. Bars represent the averages and SEM of at least three biological replicates.

**Suppl. Fig. 5.** A-B) Percentage of third (A) and fourth (B) generation mammosphere formation efficiency (% MFE) in MCF7 Empty and ETV7 cells in response to 5ng/ml IFN-β or IFN-γ calculated as number of mammospheres per well/number of cells seeded per well X 100. C) RT-qPCR analysis of the normalized expression of the genes regulated by ETV7 in MCF7 cells over-expressing ETV7 treated with 5ng/ml IFN-β (red) or IFN-γ (blue) at different time points. Relative Fold Change has been obtained normalizing the expression of the genes of interest relative to their expression in MCF7 Empty cells untreated. Bars represent the averages and standard deviation of at least three biological replicates. ** = p-value < 0.01.

**Suppl. Fig. 6.** A-B) Western Blot analysis of phosphorylated and total levels of STAT1 in MCF7 Empty and ETV7 cells in response to 5ng/ml IFN-β (A) or IFN-γ (B) at different time points. Tubulin expression was used as loading control. Blots were cropped for clarity and conciseness of the presentation. On the left of each blot is indicated the approximate observed molecular weight.

