## Supplementary figures and images for "ETV7 regulates breast cancer stem-like cell plasticity by repressing IFN-response genes"

### Supplementary Figure 1

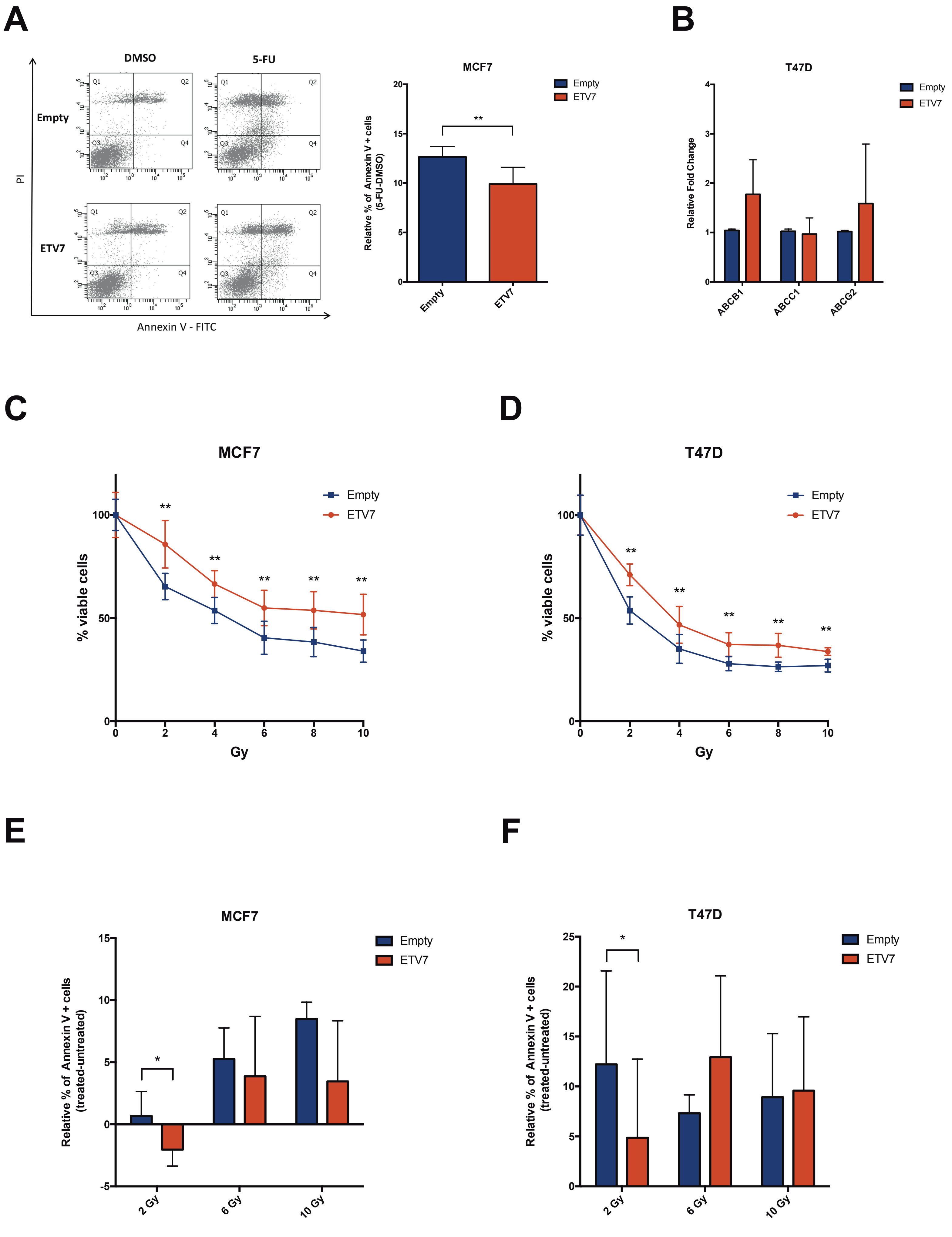

### Supplementary Figure 2

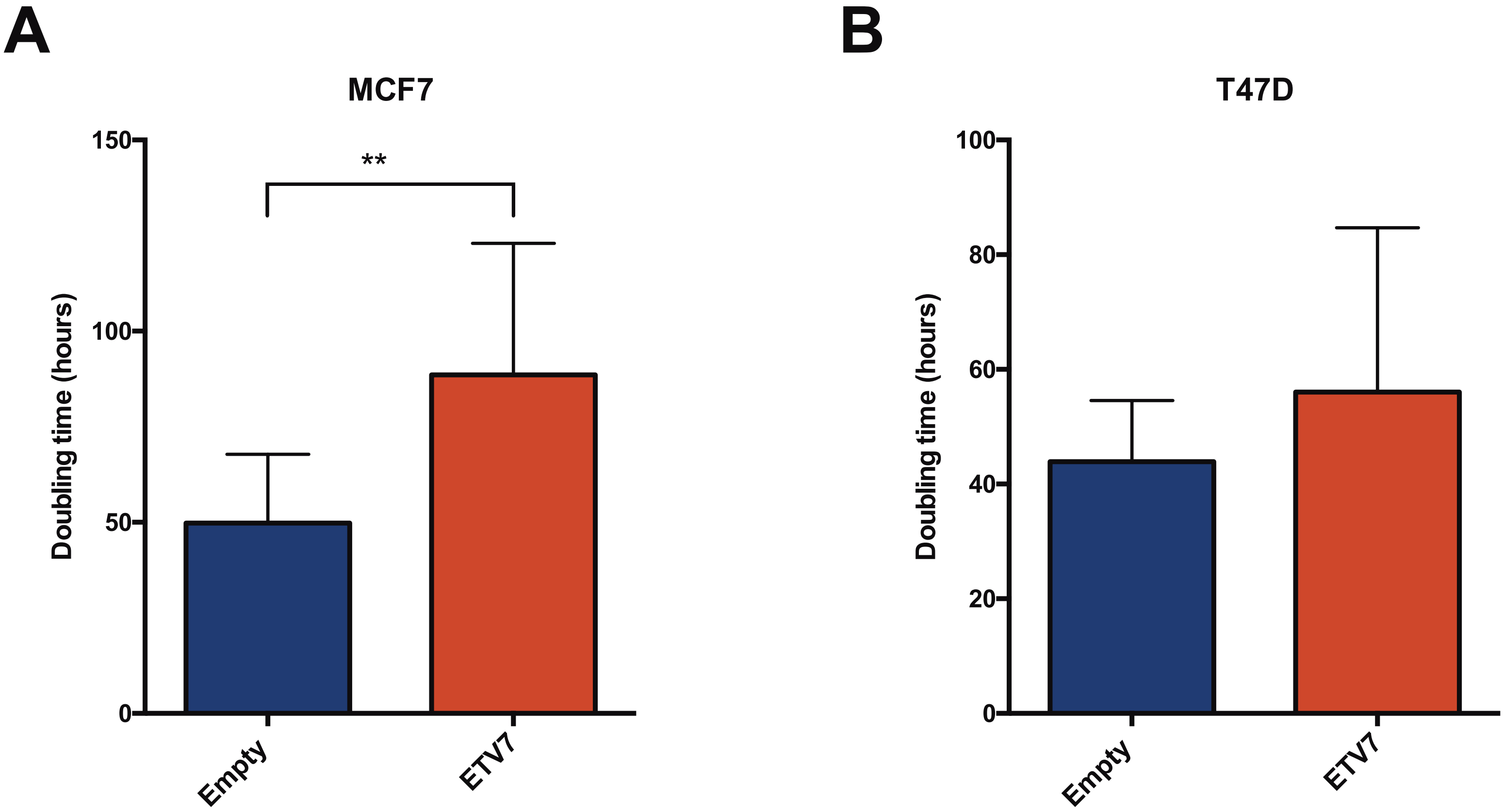

### Supplementary Figure 3

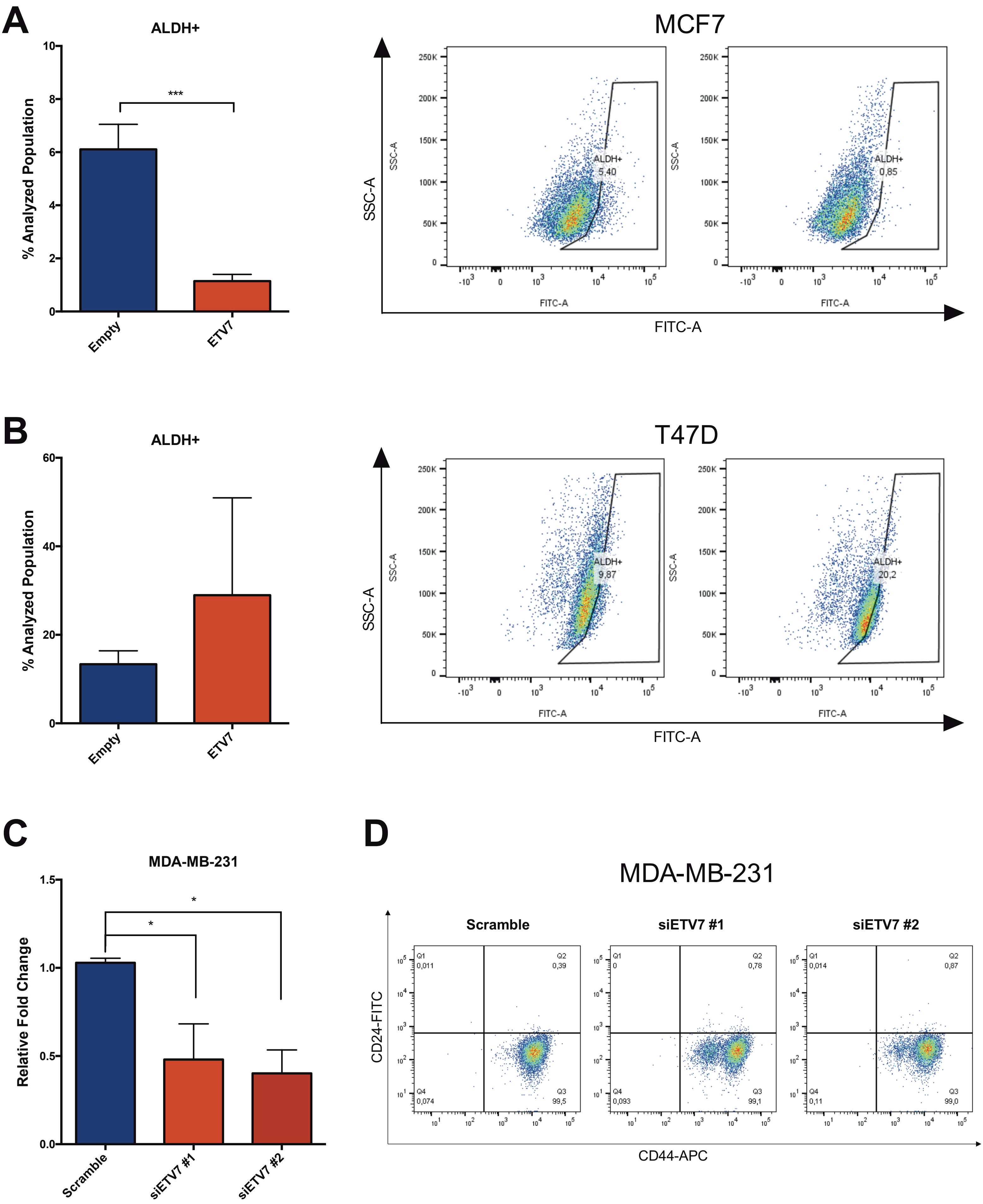

### Supplementary Figure 4

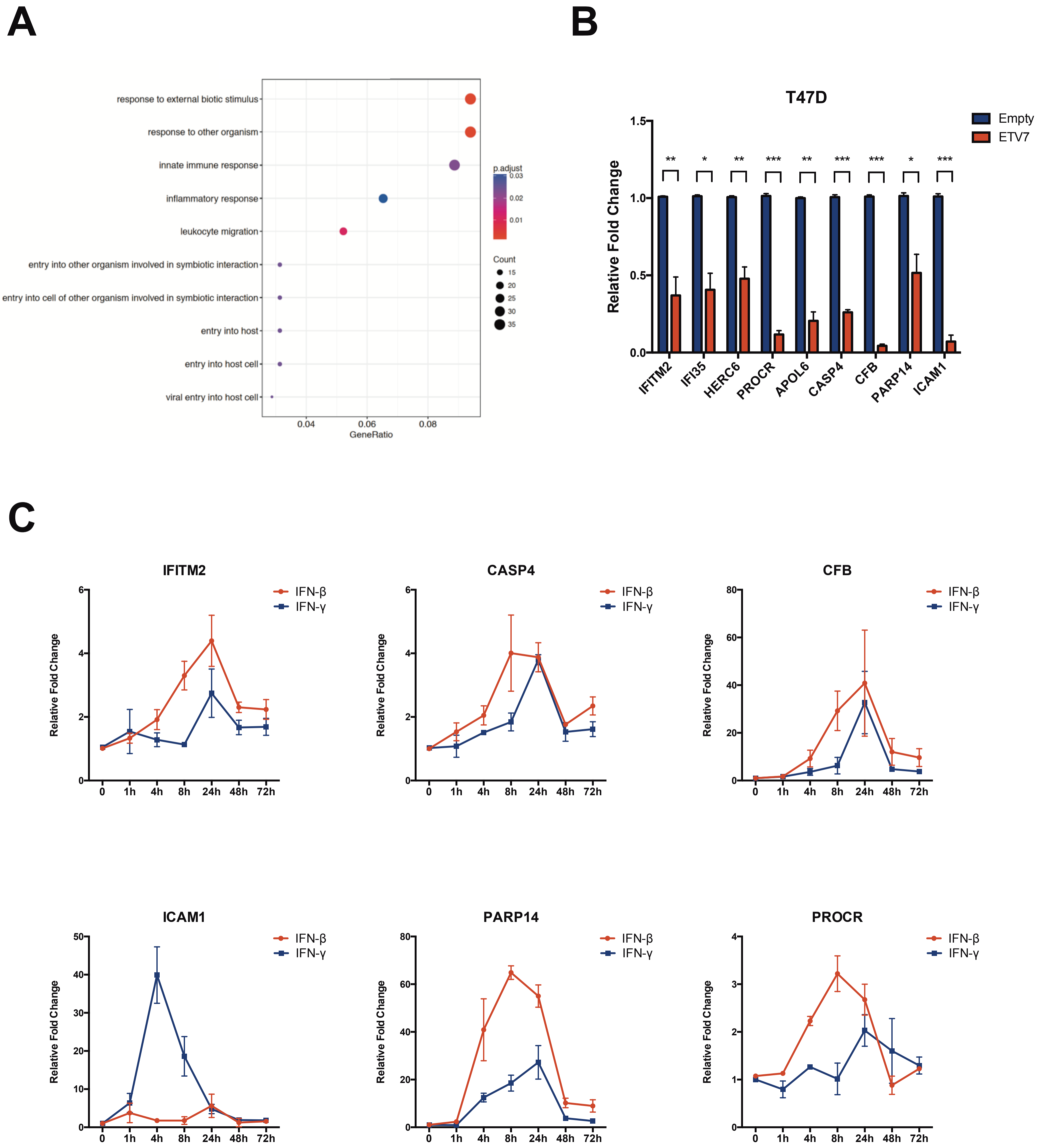

### Supplementary Figure 5

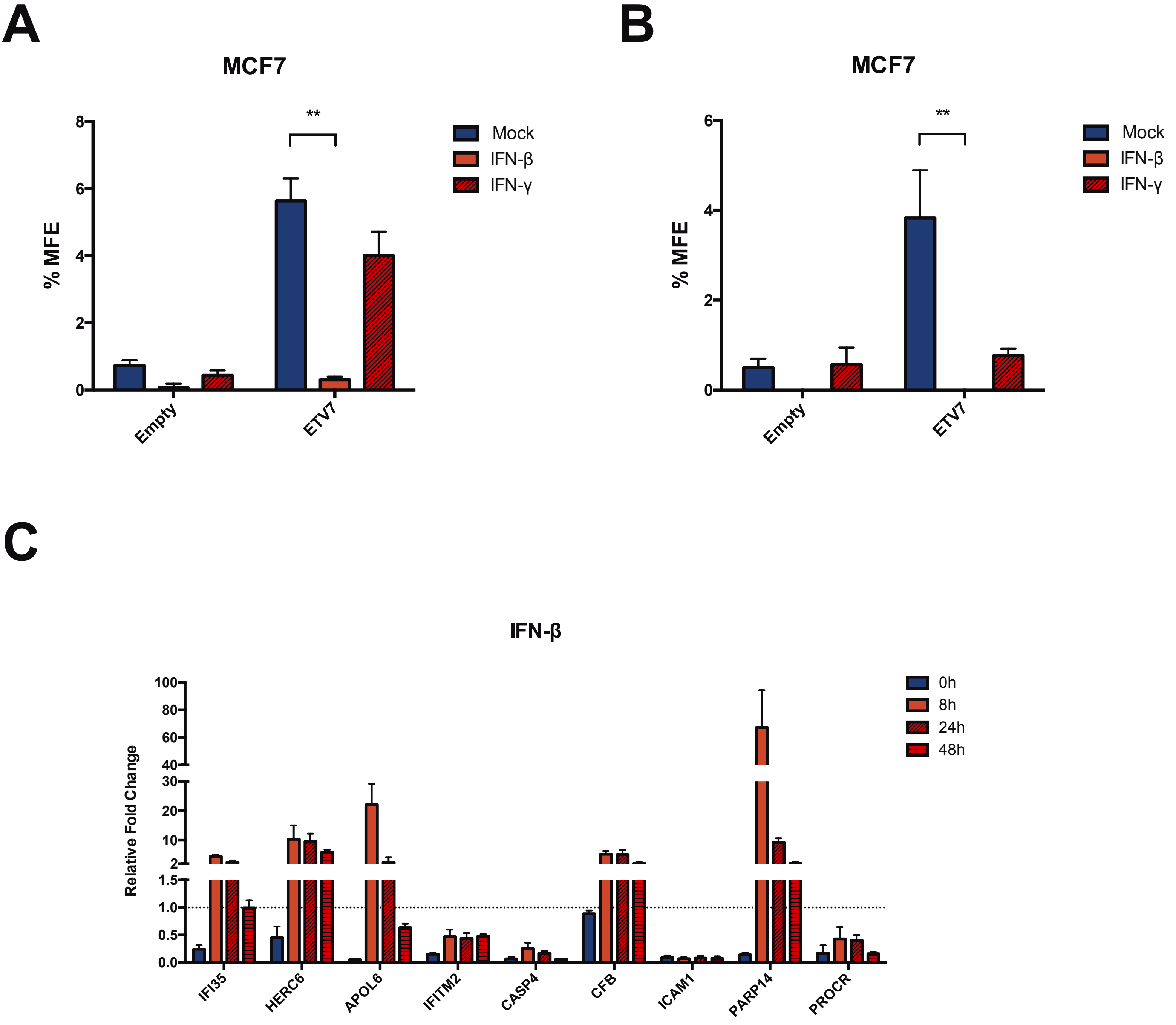

### Supplementary Figure 6

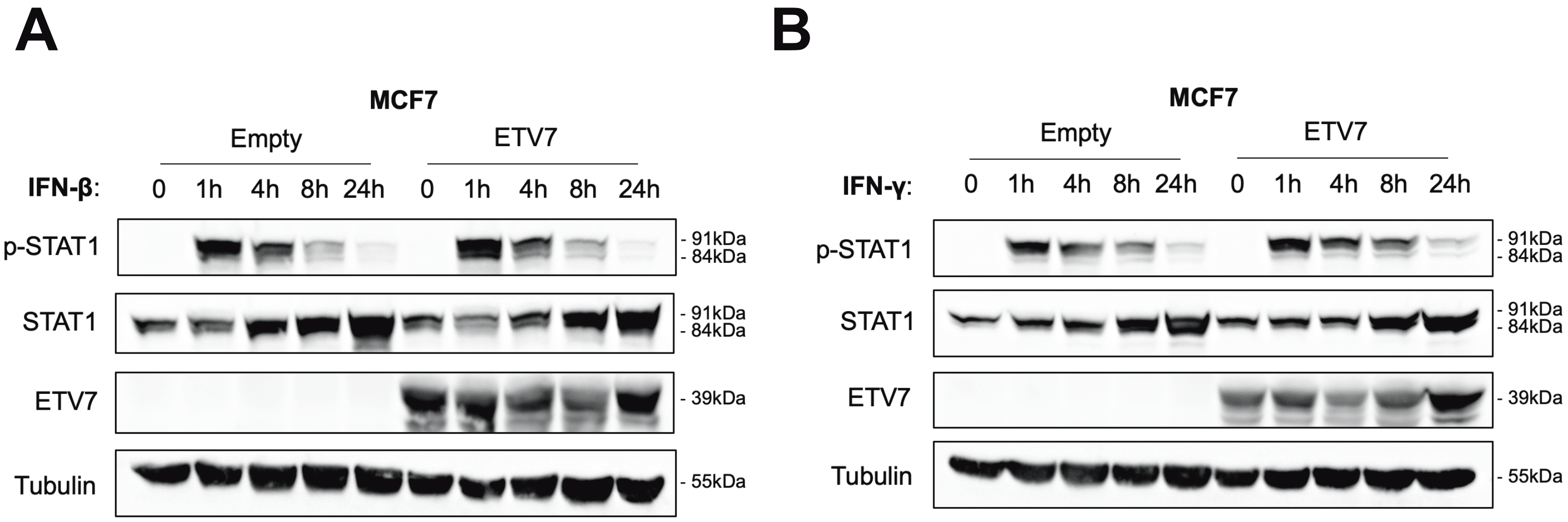
